# Daytime Colour Preference in Drosophila Depends on the Circadian Clock and TRP Channels

**DOI:** 10.1101/694315

**Authors:** Stanislav Lazopulo, Andrey Lazopulo, James Baker, Sheyum Syed

**Affiliations:** University of Miami

## Abstract

By guiding animals towards food and shelter and repelling them from potentially harmful situations, light discrimination according to colour can confer survival advantages^1,2^. Such colour-dependent behaviour may be experiential or innate. Data on innate colour preference remain controversial in mammals^3^ and limited in simpler organisms^4–7^. Here we show that when given a choice among blue, green and dim light, fruit flies exhibit an unexpectedly complex pattern of colour preference that changes with the time of day. Flies show a strong preference for green in the early morning and late afternoon, a reduced green preference at midday and a robust avoidance of blue throughout the day. Genetic manipulations reveal that the peaks in green preference require rhodopsin-based photoreceptors, and are controlled by the circadian clock. The midday reduction in green preference in favour of dim light depends on the Transient Receptor Potential (TRP) channels dTRPA1 and Pyrexia (Pyx), and is also timed by the clock. In contrast, blue avoidance is primarily mediated by class IV multidendritic neurons, requires the TRP channel Painless (Pain) and is independent of the clock. With unexpected roles for several TRP channels in *Drosophila* colour-specific phototransduction, our results reveal distinct pathways of innate colour preference that coordinate the fly’s behavioural dynamics in ambient light.

---

To address colour preference of *Drosophila*, we performed multi-day behavioural experiments providing flies with choices of space illuminated with blue, green or red light of roughly equal intensity (Fig.1a, Extended Data Fig.1a,b). Since flies do not possess red-sensitive photoreceptors^5^, the red zone in our experiments is likely perceived as a dimly lit region (Extended Data Fig.2a). Individual flies in the experiments were housed in glass tubes covered with three plastic colour filters dividing each tube into three equal colour zones with food available at one end. Day-night conditions were created by alternating 12 hours of constant light with 12 hours of complete darkness (LD). To avoid possible bias towards food, filter positions were combined in all six possible ways. A custom-written computer program quantified fly preference for colours by automatically locating individuals in their chosen colour zone every minute; these were then averaged over 1 hour time intervals, providing the distribution of flies across the three colour zones throughout the day-night cycle (Fig.1b, Extended Data Fig.3). To our surprise^4,7,8^, we found that wild-type (Canton S, CS-1) flies consistently avoided blue light during daytime hours (Fig.1b). In contrast, their preference for green changed with time: around Zeitgeber time 2 (ZT2), 2 hours after lights turned on, flies preferred to occupy their respective green colour zones (Fig.1b,c). During the middle of the day, their preference for green decreased with the fly population split roughly equally between green and dim zones. Approximately 1 hour before lights turned off, flies returned to green. This time-dependent pattern in colour preference was consistent over at least 10 days diverging only on the first day, presumably during which flies acclimate to the new LD environment. During the dark phase of LD cycles or in constant darkness (DD), flies randomly distribute themselves among the three zones, indicating that light is essential for generating the observed pattern (Extended Data Fig.2b). Multiple wild-type strains and *w*^*1118*^ with *mini-white* showed similar patterns in colour preference, suggesting the behaviour is not specific to one laboratory strain (Extended Data Fig.4). Temperature was found to be constant across colour zones and positions within the tube (Extended Data Fig.1c), thus ambient temperature differences did not drive flies’ choice of location in the tubes.

**Figure 1.**
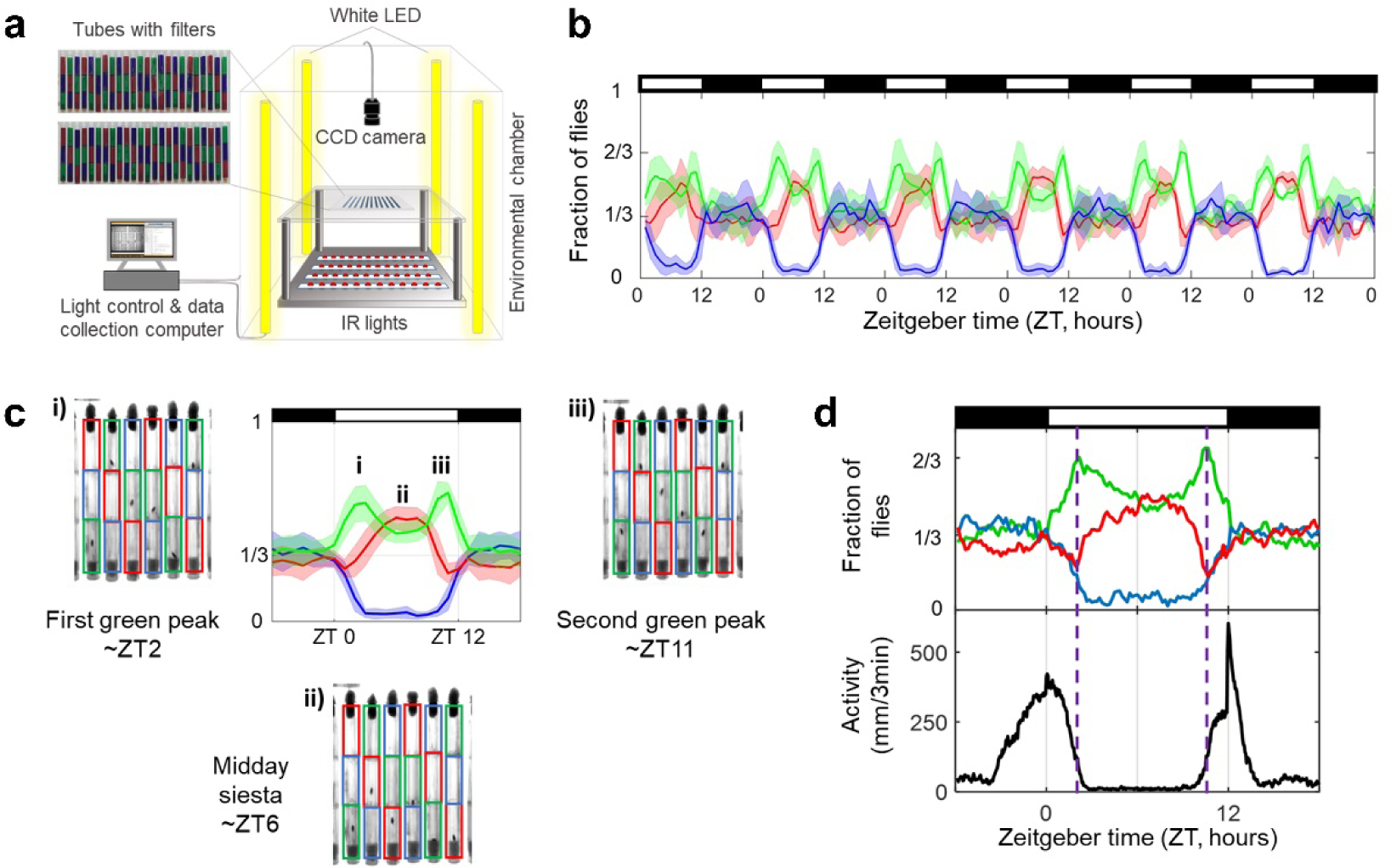
Flies exhibit a systematic change in colour preference during day. **a**, Schematic of the experimental setup. Individual fly positions in tubes divided into three colour zones are recorded every minute for multiple days using a CCD camera. Broad-band white LED lamps simulate alternating light-dark (LD) conditions while infra-red lights permit CCD to continuously track flies **b**, The distribution of flies in colour zones during 6 days of experiment under 12/12 LD conditions. The fraction of flies in green, blue and red zones is shown in the corresponding colour line. Shaded band surrounding each line represents standard deviation in data from six independent experiments. Each experiment used 18 or 24 flies (Total number N=126). **c**, Typical fly locations during green peaks and midday change in green/red preference. The coloured rectangles of corresponding filters overlay the video frames. The single-day colour preference is the average of a six-day experiment with deviations between days shown in bands. **i-iii** show single greyscale frames with indicated colour filters during the first green preference peak, the midday green/red switch and the second green peak. **d**, Correlation between colour preference (top) and locomotor activity (bottom, black line) averaged in 3 min intervals. Activity was measured as the displacement of flies between two consecutive frames and binned into 3 min intervals. Vertical dashed lines show the timing of peaks in green colour preference. **b-d**, Black and white horizontal bars indicate dark and light portions of LD cycle, respectively.

We conducted two additional experiments to query sensitivity of our observations to light intensity. In one experiment, we kept green/dim intensities constant and measured colour preference profiles for different intensities of blue light (Extended Data Fig.5a,b). Though blue avoidance decreased for lower intensities, the negative response towards blue persisted down to the lowest tested intensity. Moreover, patterns in green/dim preference remained largely unaffected throughout (Extended Data Fig.5a). In another experiment, we addressed the role of intensity in green preference through a two-choice assay, in which flies had to choose between two green zones of different intensities (Extended Data Fig.5c-e). We hypothesized that if the early-morning and late-afternoon preference for green was due to intensity rather than colour, then a choice between any two green zones where one is relatively dimmer should reproduce the pattern seen in green vs. red (Extended Data Fig.5c) by driving flies to the ‘brighter’ zone at the appropriate times. Instead, we found that only when the dimmer green was tenfold less intense than the brighter, did flies show preference patterns similar to that for green/red (Extended Data Fig.5d). In the middle of the day, however, flies always favoured the lower intensity choice (Extended Data Fig.5e). Together, these data support the view that flies prefer green and avoid blue based on wavelength rather than intensity, but shift their midday preference based on intensity.

Visual examination of the video recordings revealed that the middle of the day switch in green/dim preference was strongly affected by a heightened tendency around midday for flies to stay close to food (Fig.1c, Sup. Movie ZT4-12). However, the drive to be near food was insufficient to overcome aversion of blue light, as flies in tubes with food in the blue zone rested in the adjacent zone (green or dim) and made brief incursions into blue only to feed (Extended Data Fig.6). The strong aversion to blue seen in our paradigm would be consistent with blue light being harmful to *Drosophila*^9^. Our findings are surprising, however, given a number of prior phototaxis studies^4,7,8^ concluding that if given a choice of colours, flies invariably prefer light of the shorter wavelength. This discrepancy in fly blue light response is likely related to idiosyncrasies of the previous phototaxis assays which used low light intensities (≲ 100 μW/cm^2^) and measured a group response shortly after flies were exposed to stimuli, possibly eliciting an escape behaviour. Flies in our paradigm are provided with light of intensity closer to what they experience in nature (see Methods), monitored individually, and over multiple days. In contrast to the monotonic avoidance of blue, the two peaks in green preference resulted from flies choosing to reduce their locomotor activity and remain almost exclusively under green light (Fig.1c, Sup. Movie ZT20-4 and ZT4-12) and were timed with the conclusion of their morning (M) and the start of their evening (E) bursts in locomotor activity^10–12^ (Fig.1d). Interestingly, the presence of colour filters did not alter the heights and widths of the M and E bursts but did lower midday activity (Extended Data Fig.7a). Thus, affording flies the ability to freely choose ambient light colour seems to affect patterns in their daytime sleep and feeding behaviours^13,14^.

We next set out to identify the phototransduction pathways that regulate *Drosophila* colour preference. First, we focused on the robust avoidance of blue light. To test the role of the fly visual system, we measured colour preference in strains bearing mutations in genes that would disrupt vision: *glass* (*gl*^*60j*^), *no receptor potential A* (*norpA*^*P24*^), and a line expressing the pro-apoptotic gene *head involution defective* (*hid*) under control of the *Glass Multimer Reporter* (*GMR-hid)*. Flies homozygous for *gl*^*60j*^, a null mutation of *glass*, have defects in the development of retinal photoreceptors, ocelli and Hofbauer-Buchner (H-B) eyelets^15^; *norpA*^*P24*^ flies have a null mutation in the phospholipase C-β that disrupts the phototransduction cascade^16^; *GMR*-*hid* flies lack eyes, H-B eyelets and ocelli (Extended Data Table 1). We also measured mutants lacking only ocelli (*oc*^*01*^). Surprisingly, for all blind and ocelli-less mutants, avoidance of blue light was near wild-type level (Fig.2a,b; Extended Data Fig.8a,b). Reduction of the blue light intensity by half resulted in a lower avoidance for blind flies, but not wild-type (Extended Data Fig.8f). These data raised the possibility that both visual and non-visual receptors mediate blue avoidance, with non-visual pathway playing an important role at a higher intensity.

**Figure 2.**
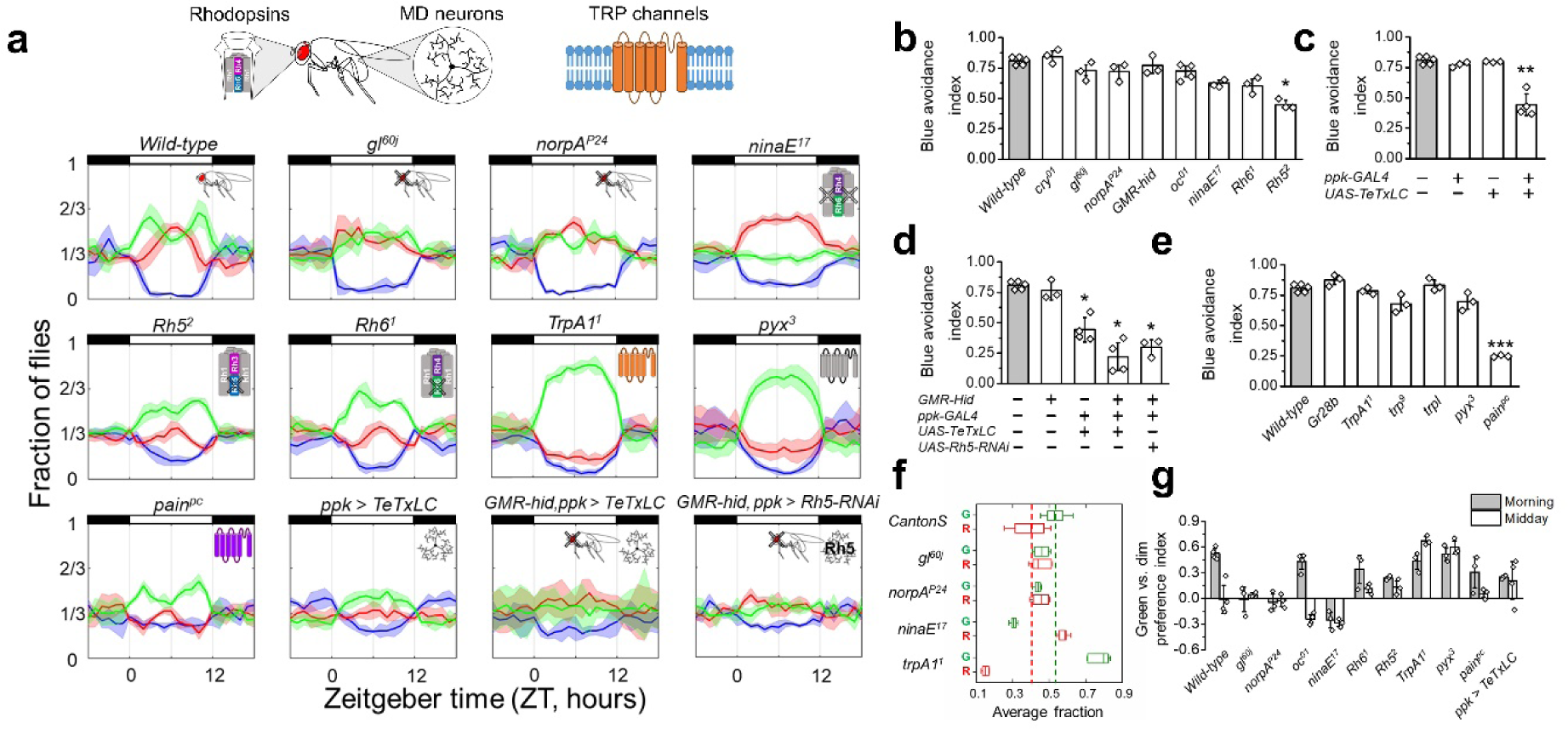
Contributions of visual photoreceptors, md neurons and TrpA channels to colour preference. **a**, Preference among blue, green and dim (red) light for each genotype during one day. Top shows affected systems of interest. The fraction of flies in green, blue and red zones shown in corresponding colour. An average 24 hr. data-set (solid lines) and associated standard deviation (s.d., shaded bands) was generated from 3-5 experiments, each conducted for 4-6 days, with 18 or 24 flies of each genotype in each experiment. Black and white horizontal bars above indicate dark and light part of LD cycle. **b-d**, Comparison of blue avoidance for flies with visual mutations (**b**), with blocked neurotransmission in multidendritic neurons (**c**), with impairment of both vision and md neurons (**d**). **e**, Comparison of blue avoidance with elimination of Gr28b and TRP channels. **b-e**, The avoidance index was calculated (see Methods) for each hour from ZT 2-10, then averaged. **f**, Average fractions of wild-type and blind flies in green and red zones during light part of the day. **g**, Preference between dim and green lights near timing of the first green peak ZT 2 and midday ZT 6. Preference for green is shown as positive and preference for dim light as negative. **b-e**,**g**, Error bars indicate s.d. between independent experiments. *P<0.05, **P<0.01, ***P<0.001; Fisher’s exact test. For exact P-values, see Extended Data Fig.8g.

In addition to rhodopsins, fruit flies possess the blue light sensitive photopigment Cryptochrome (Cry)^17^ that is involved in setting photoperiod in animals. To test whether *cry* is involved in blue avoidance, we measured a *cry* knock-out mutant *cry*^*01*^. However, no reduction of blue avoidance was observed (Fig.2b and Extended Data Fig.8c). At least three rhodopsins in the fly visual system are sensitive to blue light of wavelength ∼450 nm^18^ including Rhodopsin 1 (Rh1), which is expressed in R1-6 photoreceptor cells, Rhodopsin 2 (Rh2), which is expressed in the ocelli and Rhodopsin 5 (Rh5), which is expressed in the R8 photoreceptor cells of the compound eye and in H-B eyelets. In addition to their expression in visual photoreceptors, these Rhodopsins have been found in other cells in larvae: Rh1 is found in TrpA1-positive neurons^19^, and Rh5 is expressed in multidendritic (md) neurons^20^. To test the hypothesis that rhodopsins outside the visual system might be critical for blue avoidance, we measured flies lacking Rh1 (*NinaE*^*17*^) or Rh5 (*rh5*^*2*^) (Fig.2a). Of the two, only *rh5*^*2*^ showed significantly lower blue avoidance (Fig.2b). This result, together with wild-type response of blind and ocelli-less flies, suggests that an Rh5-dependent nonvisual pathway mediates the robust avoidance of blue.

Since Rh5 is expressed in class-IV multidendritic (md) neurons in larvae^20^ where these neurons mediate blue light avoidance^21^, we hypothesized that md neurons in adult flies might play a similar role using Rh5 as the light receptor. Consistent with that hypothesis, flies in which md signalling was blocked with *pickpocket* driven (*ppk*-*Gal4*) ^22,23^ tetanus toxin light chain (*TeTxLC*), showed a strong reduction in blue light avoidance (Fig.2a,c). Blocking md neurons signalling in blind *GMR*- *hid* flies retained the flies’ ability for locomotor activity (Extended Data Fig.7b) but led to an even stronger reduction in avoidance behaviour (Fig.2d). Consistent with our hypothesis, RNAi-mediated depletion of Rh5 in *ppk*-expressing neurons of blind flies showed a similar reduction of blue avoidance (Fig.2d). Together, these data suggest that Rh5 functions in the adult class-IV md neurons to mediate avoidance of blue light.

Gustatory receptor *Gr28b* and *trpA1* are also expressed in larval class-IV multidendritic neurons where they are necessary for proper response to blue light^21^. We wondered if larval functions of *Gr28b* and *trpA1* in light response are conserved in adult flies. Surprisingly, neither *Gr28b* nor *trpA1* mutants showed a reduction in blue avoidance in the adult (Fig.2a,e). To determine if a different member of the TRP family could be involved, we broadened our studies to flies with a null mutation in one of the other related channels: *trp*, and *trpl* which are required for classical phototransduction, *painless* which mediates multiple noxious stimuli and *pyrexia*, which is mainly responsible for temperature nociception^24,25^. Out of all tested mutants, only *painless* showed a reduction in blue avoidance (Fig.2a,e). Painless has been previously implicated in aversive responses to noxious heat, mechanical and chemical stimuli but not bright light^25^. To our knowledge, the present study is the first demonstration of a critical function for *painless* in *Drosophila* phototactic behaviour.

Although deficiencies in the visual pathway (*gl*^*60j*^, *norpA*^*P24*^, *GMR*-*hid, trp*) did not affect blue light avoidance, the mutations eliminated preference for green, resulting in an equal number of flies in green and dim regions (Fig.2a,f,g; Extended Data Fig.3b, right). Two rhodopsins in *Drosophila* are sensitive to green light: broadly sensitive Rh1 that is expressed in photoreceptors R1-6 and thought to be involved in dim light vision and green-specific Rh6 that is restrictively expressed in photoreceptor R8 and involved in colour-specific vision^5,18,26^. When tested, the *rh6*^*1*^ mutant showed near wild-type peaks in green preference and only slightly reduced midday green preference (Fig.2a,g). In comparison, flies lacking the Rh1 (*ninaE*^*17*^) showed profound changes, completely losing preference for green (Fig.2a,g) and dwelling in dim light throughout the day (Fig.2f). These results suggest that while Rh6 mediates some green light information for the behaviour, Rh1 is primarily responsible for eliciting preference for green light.

Consistent with a model in which green preference is controlled independently of blue avoidance, *rh5*^*2*^ and *painless* mutants with reduced avoidance of blue, still show the two peaks in green and the midday switch in green/dim preference (Fig.2a,g). Conversely, the *trpA1* and *pyrexia* mutants that showed normal blue light avoidance, lost the midday green/dim switch and instead, preferred green most of the day (Fig.2a,g,f). TrpA1 and Pyrexia (Pyx) have been primarily described as molecular heat sensors in *Drosophila* nociception^24,25^. However, they were also shown to function in other behaviours at temperatures far from their direct activation range, with TrpA1 controlling midday rest and activity^13,27^ and Pyx mediating temperature-entrainment of the circadian clock^28^. Given their newly uncovered roles in colour preference, TrpA1 and Pyx, like rhodopsins^19,20^, appear to play roles in both photo- and thermo-driven behaviours.

The genetic manipulations discussed thus far can be categorized into two groups according to the phenotypic changes they produce: one group, like *painless*, that clearly separate blue avoidance from green preference; and a second group, like disruption of md neuron function, that produce related changes in both behaviours (Fig.2a). The two conflicting models of avoidance-preference independence likely result from the design of our three-choice assay in which change in one behaviour (avoidance or preference) can sometimes produce a change in the other. In order to resolve differences between the divergent models, we tested flies in two-colour choice assays with either blue and red or green and red (Fig.3a,c). Wild-type flies showed blue avoidance and green/dim switch in two-colour assays (Fig.3b,d) similar to those observed in three-colour assays (Fig.2b,g). Since blue avoidance does not require green and the green/dim switch occurs even without blue, the two behaviours are indeed independent. Consistent with this model of separate avoidance and preference pathways, *painless* flies showed defective avoidance of blue (Fig.3a,b) but near wild-type patterns in green/dim preferences (Fig.3c,d) while *gl*^*60j*^, *norpA*^*P24*^, *ninaE*^*17*^, *TrpA1*^*1*^ and *pyx*^*3*^ mutants showed wild-type avoidance of blue but atypical behaviours in green vs. red assays. In contrast, *Rh5*^2^ and *Rh6*^*1*^ flies exhibited simultaneous changes in blue avoidance and green preference in the three-colour assay. The two-colour experiments clarified that changes in *Rh5*^2^ green preference and *Rh6*^*1*^ blue avoidance are not significant (Fig.3). Simultaneous changes appear in the three-colour assay possibly because the mutations affect co-expression of Rh5 and Rh6^20^. Lastly, *ppk*>*TeTxLC* flies also showed simultaneous changes in the three-colour assay and the two-colour assays appeared to validate those results by showing decreased avoidance of blue and the absence of the midday green/dim switch (Fig.3). However, while both controls showed wild-type blue avoidance, *UAS-TeTxLC* flies already had an abnormal green preference (Extended Data Fig.8e), suggesting that *ppk*>*TeTxLC* changes in green/dim behaviour were not due to md neurons. Thus, the two-colour assays further substantiate results from the three-colour paradigm supporting a separation of the aversion and preference pathways.

**Figure 3.**
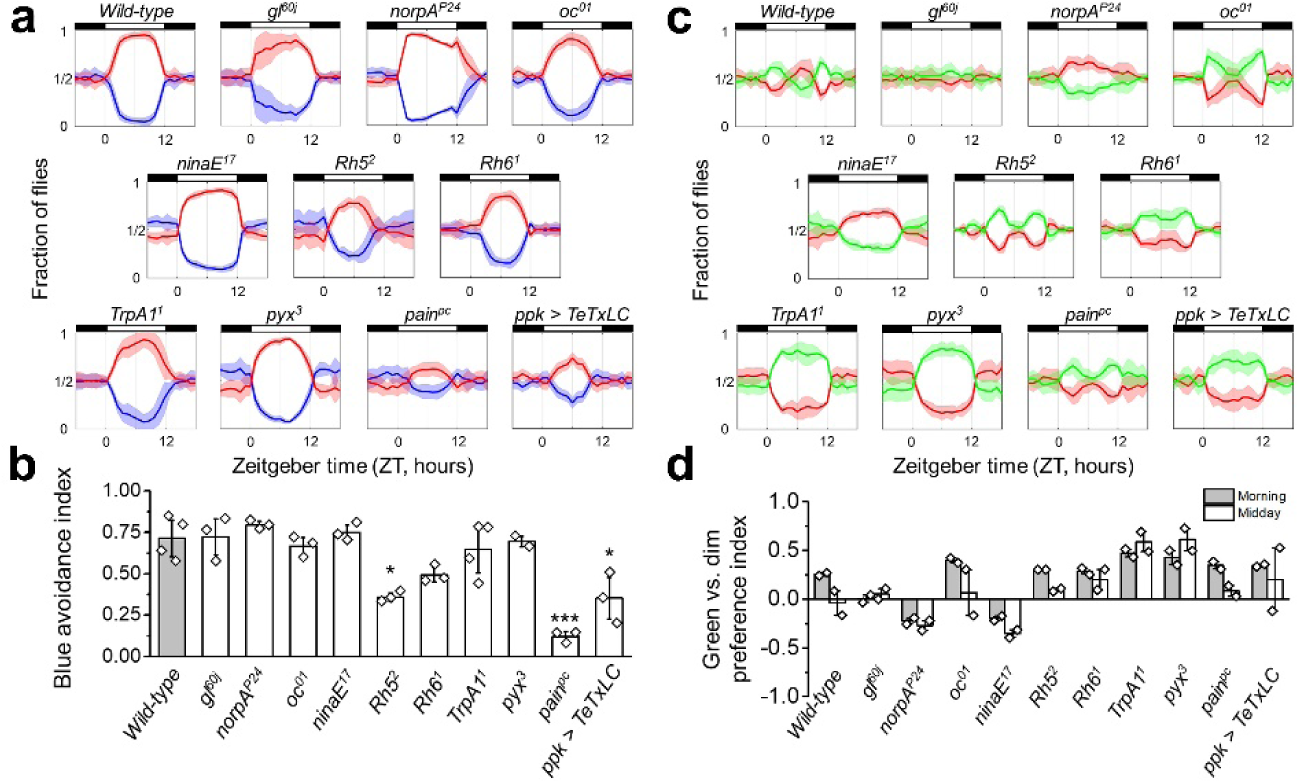
Colour preference in two-choice assays. **a**, Average daily preference between blue and dim light for each genotype. The fraction of flies in blue and red (dim) zones shown in solid line of the corresponding colour and standard deviation is shown in shaded band around each line. Average preference is calculated from 2-3 repeats of 4-6 day-long experiments with 18 or 24 flies of each genotype. Black and white horizontal bars above indicate the dark and light parts of the LD cycle. **b**, Comparison of blue avoidance between different genotypes. The avoidance index was calculated (see Methods) for each hour from ZT 2-10, then averaged. **c**, Average daily preference between green and dim lights for each genotype. Apart from the difference in colour choices, all experimental details are the same as in a. **d**, Preference between green and dim around ∼ZT 2 and around midday ∼ZT 6. Preference for green shown as positive and preference for dim light as negative. **b, d**, Error bars indicate standard deviation between independent experiments. *P<0.05, **P<0.01, ***P<0.001; Fisher’s exact test. For exact P-values, see Extended Data Fig.8h.

Several lines of evidence support a role of the circadian clock in fly colour preference: the periodic pattern in choice behaviour, the loss of periodicity in constant light (Extended Data Fig.2c) and appearance of the second green peak in apparent anticipation of the E burst in activity (Fig.1d). To further elucidate the involvement of the clock, we measured colour preferences in flies carrying mutations in the obligate clock gene *period* (*per*). Previous studies^29^ have identified mutants with short (*per*^*S*^), long (*per*^*L*^) and abolished (*per*^*0*^) circadian rhythms. We exposed wild-type and clock mutants to 19, 24 and 29.5 hour LD cycles, which correspond to the *per*^*S*^, wild-type and *per*^*L*^ endogenous rhythms, respectively (Fig.4a-c and Extended Data Fig.9). Flies placed in external LD cycle matching their innate rhythm showed wild-type behaviour with two peaks and a midday switch in green/dim preference (Fig.4a-c, along top-left to bottom-right diagonal). The first peak in green closely followed the end of the M burst in activity and the second peak preceded the start of the E burst in activity (Extended Data Fig.9a). Consistent with a clock-controlled modulation of colour preference, flies with a non-functional clock showed a monotonic preference for green regardless of the length of the day (Fig.4d and Extended Data Fig.8d). However, when rhythmic flies were placed in external cycle differing from their innate rhythm, their colour preference behaviour was modified. When the internal clock ran faster than the external cycle, such as *per*^*S*^ exposed to 24 or 29.5 hour photoperiod, the M burst occurred in the dark phase (Fig.4a, black arrows and Extended Data Figs.9), and the first peak in green preference was absent. However, when the external cycle was faster and the M activity was initiated by lights-on, such as wild-type in 19 hours or *per*^*L*^ in 24 hours photoperiod, the first green peak followed within the next ∼2.5 hours (Fig.4b,c). As a result of this interplay between the internal and external cycles, the first green peak, when present, always appeared around ZT2.5 or ∼2.5 hours after lights-on (Fig.4e, open symbols). Similarly, the second green peak was absent when the E activity occurred in the dark phase, as in the case of wild-type in 19 hour or *per*^*L*^ in 24 hour photoperiod (Fig.4b,c, white arrows and Extended Data Figs.9). In conditions where the E burst in activity occurred in the light phase, for instance wild-type flies in 24 hour or 29.5 hour cycle (Fig.4b, white arrows), the second green peak moved in ZT concomitant with shifts in timing of the evening locomotor activity, but always appearing ∼1 hour before the E burst. Consequently, the second green peak, when present, appeared at a different ZT depending on relative paces of the internal and external cycles (Fig.4e, filled symbols). While these data hint at a strong association between peaks in green preference and bursts in locomotor activity, comparison between *per*^*0*^ and *TrpA1*^*1*^ data suggest that the features in colour preference and locomotion may be regulated independently. While both TrpA1 and a functional clock are needed for the midday switch in green preference (Fig.2a and Fig.4d), only the clock, still functional in *TrpA1*^*1*^ flies, is needed for the anticipatory M and E bursts (Extended Data Fig.7c).

**Figure 4.**
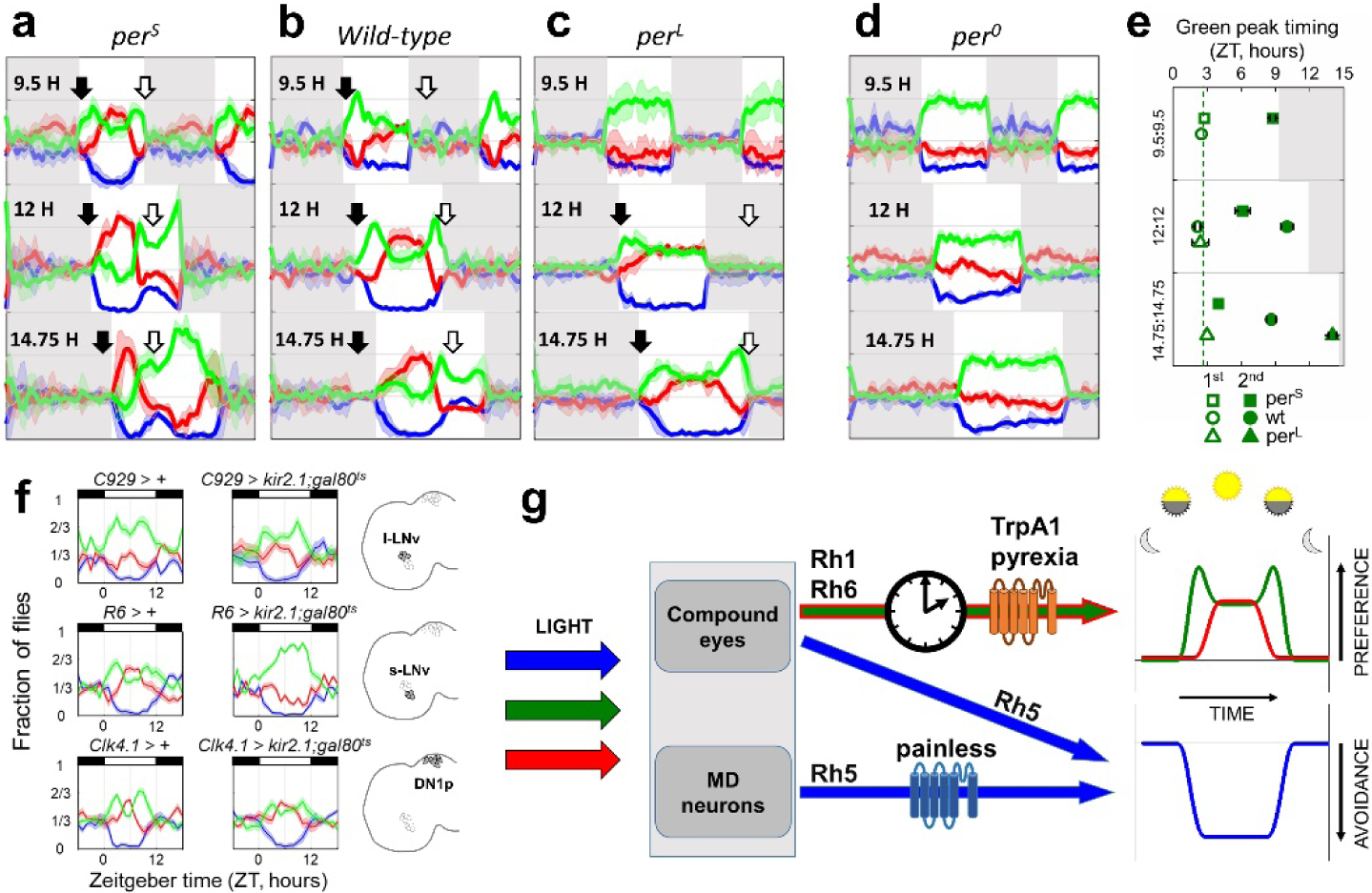
Circadian clock regulates the timing of changes in preference between green and red colours. **a-d**, Wild-type and clock mutants in three different LD cycles (9.5:9.5 LD, 12:12 LD, 14.75:14.75 LD). Grey area shows dark part of LD cycle; black and white arrows indicate the timing of morning and evening bursts in locomotor activities respectively (for colour preference with activity see Extended Data Fig.7). **e**, Timings of the first (open symbols) and the second (filled symbols) green peaks in Zeitgeber time. Peaks for *per*^*S*^ shown with squares, wild-type with circles and *per*^*L*^ with triangles. **f**, Silencing s-LNvs leads to loss of green/dim switch in preference. Comparison of colour preference of flies with silenced l-LNvs (top, C929-Gal4), s-LNvs (middle, R6-Gal4) and DN1ps (bottom, Clk4.1-Gal4) and corresponding control flies. Targeted neurons are shown on the right. **a-d, f**, The fraction of flies in green, blue and red (dim) zones shown in corresponding colour, standard deviation between two different experiments is shown in the shaded bands; 18 or 24 flies were used for each experiment. Black and white horizontal bars indicate dark and light parts of LD cycle. **g**, Model of the receptive and transduction pathways contributing to the colour preference behaviour.

To identify which of the ∼150 clock cells in the fly brain are primarily responsible for conferring rhythmicity to colour preference, we used the *UAS*-*Gal4* system to specifically target the leaky inward rectifier K+ channel (*UAS-kir2.1*) to silence large (l-LN_v_, with *C929-Gal4*) and small ventral lateral neurons (s-LN_v_, with *R6-Gal4*) and a group of dorsal neurons (DN1_p_, with *Clk4.1-Gal4*), neuronal subsets important for generating the daily M and E activity bursts^10,11^. We used the temperature dependent repressor of *Gal4, tub-Gal80*^*ts*^ to raise animals under restrictive conditions and silence clock neurons only in the adult stages. The experiments revealed that silencing with *R6-Gal4* produces patterns (Fig.4f) that most closely phenocopies circadian arrhythmic flies (Fig.4d and Extended Data Fig.8d). In contrast, *C929-Gal4* and *Clk4.1-Gal4* driven silencing largely retained wild-type behaviour. This result suggests that proper electrical firing of the small ventral lateral neurons is critical for the midday switch in colour preference and generation of the twin peaks in green preference.

Here we show that *Drosophila* have a heretofore undescribed preference for colour that varies with the time of day. The circadian variation in *Drosophila* raises the intriguing possibility that colour preference in humans may also depend on the time of day, in addition to its dependence on seasons^30^. Results from our study further suggest that green colour input is received primarily by the visual system and is strongly dependent on Rh1, TrpA1 and Pyrexia (Fig.4g). The change in preference during the day is dependent on the core clock gene *per* and relies on the activity of the small ventral lateral neurons which regulate rhythmic behaviour downstream of the circadian clock. By contrast, blue light avoidance is clock independent, detected by Rh5 in the md neurons of the body wall and requires the function of TRP channel Painless in those same neurons. The time-of-day dependent choices flies make in the ambient colour environment and the resulting alterations in their daily activity, feeding and rest, may well constitute a genetically tractable example of behaviour influencing physiology^31^. In mammals, light is known to affect mood and regulate many physiological processes, such as circadian entrainment and sleep-wake cycles. The surprising interplay between systems, genes and body parts required for colour preference in the fruit fly suggest that *Drosophila* can serve as a useful system in understanding how spectral composition of light might shape physiological and behavioural processes in higher animals.

## Acknowledgments

We are grateful to C. Desplan, J. Giebultowicz, F. Hamada, W. Joiner, M. Klein, Y. Xiang, M. Young, and the Bloomington Drosophila Stock Centre for fly stocks; J. Dallman, M. Dallman, B. de Bivort, M. Klein and J. Truman for critical comments on the manuscript; J. Dallman, M. Klein and W. Li for discussions. This work was supported in part by the National Science Foundation under grant IOS-1656603 to S.S.

## Author contribution

S.L. and S.S. conceived and designed the study, S.L and A.L performed the experiments and analysed the data, S.L. and S.S. wrote the paper with feedback from all authors, S.S. and J.B. supervised the study.

## Author Information

The authors declare no competing financial interests. Readers are welcome to comment on the online version of the paper. Correspondence and requests for materials should be addressed to S.S.

## Methods

### Fly strains

A standard laboratory strain Canton-S (CS-1) was used as the wild-type control. Two additional wild-type strains (another Canton-S (CS-2) and Cambridge B) and *w*^*1118*^ with *mini-white* were behaviourally indistinguishable from CS-1 (Extended Data Fig.4). The following lines were used in the study: *trpA1*^*1*^*(#36342), trp*^*9*^*(#9046), trpl(#29134), gl*^*60j*^*(#509), oc*^*1*^*(#24)* and *norpA*^*P24*^*(#9048)* (from Bloomington Drosophila Stock Centre); *ninaE*^*17*^, *Rh5*^*2*^, and *Rh6*^*1*^ (from Claude Desplan); CS-1, *per*^*S*^, *per*^*L*^, *R6-Gal4, Clk4.1-Gal4, C929-Gal4, pdf-Gal4, tub-Gal80*^*ts*^ and *cry*^*01* 32^ (from Michael Young); CS-2, *per*^*0*^, *cyc*^*0*^ and *tim*^*0*^ (from Jadwiga Giebultowicz); *pyx*^*3*^ (from Fumika Hamada); *GMR-Hid, Dmel/Mi{ET1}Gr28b*^*MB03888*^ and *ppk-Gal4* (from Yang Xiang); Cambridge B (CamB, from a single female caught with fruit-baited trap in Cambridge, Massachusetts, USA) and *UAS-TeTxLC* (from Mason Klein); *UAS-kir2.1* (from William Joiner). Unless otherwise noted, flies were reared on standard cornmeal/agar/yeast/molasses medium at room temperature (24°C) in ambient laboratory light. All flies used in this study are on the Canton-S or *w*^*1118*^ with *mini-white* background, all mutants are homozygous viable and previously characterized in different contexts by other authors.

### Colour preference assay

Male flies of age 2-7 days were placed in ∼6 cm long, ∼5 mm wide individual glass tubes (TriKinetics Inc.) with food and cotton plug at opposite ends, wrapped in two or three plastic colour filters. To avoid any effect of food and cotton, the filters were used in all possible combinations (two for two colours and six for three colours, Extended Data Fig.10a,b). For simplicity, tubes were arranged in order as in Extended Data Fig.10a, however, randomized ordering showed same results (Extended Data Fig.10c,d); sample sizes were chosen to reliably measure experimental parameters above noise levels. Flies were visualized with an infra-red array (LEDLightsWorld Ltd., # HK-F5050IR30-X) and recorded using a CCD camera (The Imaging Source LLC) fitted with a near-IR long-pass filter (Midwest Optical Systems Inc., LP780) under 12:12 hours light/dark cycle at 25°C and 75% RH. For Fig.4a-d and Extended Data Fig.9, experiments were also performed under 9.5:9.5 and 14.75:14.75 hours light/dark cycles. Images were captured at 10 frames/second and queried at 1-minute intervals to locate flies within colour zones (see Fly Tracking). The location data yielding the number of flies within each colour were next averaged over 1-hour intervals to visualize temporal patterns in colour preference of the population. Flies on the first, and occasionally second, day of measurement did not show a consistent pattern of colour preference and were ignored in statistical analyses. Experiments were run for 4-10 days. In the three colour assay, if 1/3 fraction of flies was positioned in a colour zone, the preference was concluded to be neutral. For values from 0 to 1/3 the colour was assumed to be avoided, for values from 1/3 to 1 colour was assumed to be preferred. For blue avoidance in three-colour-choice assay (Fig.2 b-e), the fraction of flies in blue zone was compared to the fraction of flies in the other two colours combined, according to 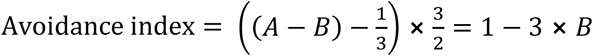, where *A* is the total fraction of flies in green and red zones and *B* is the fraction of flies in blue zone. The simplified expression with only *B* follows from the constraint *A* + *B* = 1, at all times for these three-colour experiments. Defined this way, the Avoidance index = 1 if there are no flies in blue, and 0 if 1/3 of the flies are in blue. The avoidance index was calculated for each hour from ZT 2-10, then averaged. When only two colours were involved (Fig.2g and Fig.3 b,d, Extended Data Fig.2a,5d), the standard Preference/Avoidance index = (Fraction of flies in colour A – Fraction of flies in colour B) was used.

### Fly tracking

Flies were recorded at 10 frames/second and their locations detected every second using background subtraction (Extended Data Fig.3c). After filtering out noise, only closed objects corresponding to flies remained in the binary image (Extended Data Fig.3d). To validate the method, 50 random frames from every experiment were analysed by eye and fly positions compared to the ones from computer tracking. No discrepancy was found between manual and automatic fly tracking. For colour preference, user-defined binary masks corresponding to each colour filter were next applied to background-subtracted images to determine fly position relative to the colour zones. Finally, the number of flies in each colour was divided by the total number of flies to calculate fractions at each time point. These data were typically averaged over 1-hour interval to produce the time-dependent patterns in colour preference (for example, Figs. 1b,c). Extended Data Fig.6 shows fly centre-of-mass coordinates along the tube every second. To determine activity (Fig.1d and Extended Data Fig.7,9a), displacement was calculated from the difference in fly centre-of-mass coordinates in consecutive frames and summed over 3-minute intervals.

### Statistical analysis

Videos were processed and analysed using custom-written code in MATLAB (MathWorks Inc.). The sample size was chosen based on initial experiments to ensure statistical power > 80% in finding a significant difference between different groups. First day of measurement of colour preference was excluded due to time taken by fly circadian clocks to be entrained to the external light-dark conditions. Flies within the same genotype were randomly picked for experimental groups. Investigators were not blinded to the group allocation, as each experiment requires certain genotypes for experimental and control groups. Data analysis software was blinded when assessing the results and all data collection and analysis was performed automatically by the software.

For Fig.1b, where multiple days are shown, and Fig.4f, where only one day was counted, the lines and error bands represent mean ± standard deviation between 6 (Fig.1b) and 2 (Fig.4f) separate experiments. For Fig.1c, 2a, 3a,c, 4a-d and Extended Data Figs.2,4,5,8a-f, flies were measured for 6-10 days in 2-5 separate experiments with new animals in each experiment. Lines and error bands represent mean ± standard deviation between 4-6 consecutive days, starting from the third day in each experiment. For blue light avoidance (Fig.2b-e), the Avoidance index was averaged over the “day” part of the light cycle from 2 hours after lights on to 2 hours before lights off and shown as a mean ± standard deviation from 3-5 different experiments. The small sample size of experiments (N=3 experiments, on average) did not allow the use of standard tests for normality and significance. Instead, a Fisher’s exact test with a total number of flies from all experiments was used to calculate statistical significance of differences between Preference/Avoidance indices (Extended Data Fig.8g,h).

### Locomotor activity

Fly locomotion was measured using the same video recording setup as described above, with 1-second frame rate under 9.5:9.5, 12:12, or 14.75:14.75 light/dark cycle. The activity was calculated as the distance travelled between two consecutive frames and summed over 3-minute intervals. For comparison to locomotion with colour filters, fly locomotion without filters was recorded in clear tubes under 590 μW/cm^2^ (∼2000 lux) illumination comparable to the average light intensity under colour filters (see Light properties).

### Silencing neurons

Silencing specific groups of clock neurons was performed via *UAS*-*Gal4* system driving expression of the leaky inward rectifier K+ channel (*UAS-kir2.1*)^33^ in them. *UAS-Kir2.1/R6-Gal4, UAS-Kir2.1/C929-Gal4*, and *UAS-Kir2.1/+;Clk4.1-Gal4/+* flies did not survive to adulthood. Flies with UAS-GAL4 constructs with a temperature-dependent repressor of GAL4, tubGAL80ts, were raised at 18°C to restrict the activity of GAL4 drivers during development. 1 to 5 days old male test flies and controls were transferred to 29°C for two days right before the experiment to deplete tubGAL80ts, then measured under 25°C.

### Light properties

Broad-spectrum white light (see Fig.1a and Extended Data Fig.1a,b) illumination with variable power up to 2480 μW/cm^2^ (∼8800 lux) was provided by eight symmetrically placed LED lamps (LED-88020-120V Neptun Light Inc., IL) and computer-controlled using electronics described previously^34^. Our maximum light intensity of 2480 μW/cm^2^ is less than ∼6853 μW/cm^2^ (excluding infra-red) of indirect sunlight measured around noon at 25°43’26.2 N 80°16’48.1 W. Three plastic colour filters (Roscolux #389, #58 and #19, Rosco) were used to transmit only green, blue or red light, with maximum transmittance at 528 nm, 450 nm, and 620 nm, respectively. The irradiance after filters was 626 μW/cm^2^ for green, 572 μW/cm^2^ for blue and 571 μW/cm^2^ for red (Extended Data Fig.1a,b). Average irradiance transmitted through the filters was 590 μW/cm^2^, which is comparable to the irradiance of white LED light with an illuminance of ∼2000 lux (Extended Data Table 2). For full transmission spectra data, measured with a USB-4000 (Ocean Optics) spectrometer, see Extended Data Fig.1b.

### Temperature measurement

Temperature inside the glass tubes under different colour filters and in different locations were compared and were not different from the temperature inside the incubator (25.0±0.1°C, Extended Data Fig.1c,d). Measurements were performed with a thermocouple thermometer (FLUKE 52II, FLUKE Corporation) in the dark and after 10 continuous hours of illumination.

### Data and code availability

The data that support the findings of this study are available from the corresponding author upon request. The custom computer code used in this study is available in GitHub repository (https://github.com/StasLaz/Tracking-flies).

## Extended Data Figure Captions

**Extended Data Figure 1. Properties of unfiltered and filtered light, and temperature inside the tubes. a**, Measured values for the broadband ‘white’ LED light of ∼8800 lux used in the experiments and the corresponding total irradiance of the light before and after passing through the filters. **b**, Spectral irradiance of the LED light and light transmitted through the green, blue and red filters. Peaks for the filtered light are at 450 nm (blue), 528 nm (green), and 620 nm (red).**c**, Average temperature in the green, blue, and red zones, regardless of proximity to food or cotton plug. **d**, Average temperature near food, near cotton and in the middle, regardless of filter positions. **c**,**d**, Measurements were taken with lights turned off (black bars) and after lights have been on for 10 hours (white bars). Error bars are standard deviations from 12 independent measurements.

**Extended Data Figure 2. Flies can differentiate between dim light and dark, and lose preference pattern in constant conditions. a**, Average daily preference between red and dark zones for wild-type flies. The dark zone was obtained by overlaying three colour filters (Rosco #27, #382, #389). The fraction of flies in red and dark zones shown in red and black. Standard deviation between 4 consecutive days is shown in shaded bands around each line. 20 flies were used for the experiment. **b**,**c**, Distribution of flies across colour zones in 12:12 LD conditions, followed by constant darkness (**b**) or constant light (**c**) (N=24, each experiment). The fraction of flies in green, blue and red zones shown in the corresponding colour. **a**,**b**, Black and white horizontal bars indicate dark and light conditions..

**Extended Data Figure 3. Experimental setup with tubes in filters. a**, Typical spatial arrangement of tubes (each approximately 6 cm long and 5 mm wide) in an experiment with three colour filters (green, blue, red) and two colour filters (green, red); N=132. **b**, Example frames of automatic tracking of flies in tubes with filters (right) and without filters (left). **c**, Background subtraction was applied after stabilizing video. **d**, Flies (green objects) were detected following digital filtering and noise elimination. **e**, Example of 12 randomly picked single fly trajectories, moving between green (G), blue (B) and red (R) zones over 24 hours. Fly positions were determined every minute but data shown are 5 minutes apart for clarity. **f**, Population-averaged colour preference of the 12 flies. As with the individual data, the time interval here is 5 minutes. **g**, Same colour preference but averaged over 1-hour intervals.

**Extended Data Figure 4. Preferences for colour in additional wild-type genotypes. a**, Preference among three zones for each genotype during one day. The fraction of flies in green, blue and red (dim) zones shown in the corresponding colour. An average 24 hour data (solid lines) and associated standard deviation (s.d., shaded bands) were generated from 3 experiments, each conducted for 4-6 days, with 18 or 24 flies of each genotype in each experiment. Black and white bars indicate dark and light part of LD cycle. **b**, Comparison of blue avoidance. The avoidance index was calculated (see Methods) for each hour from ZT 2-10, then averaged. **c**, Preference between green and dim light near timing of the first green peak ZT 2 and midday ZT 6. Preference for green is shown as positive and preference for dim light as negative.

**Extended Data Figure 5. Colour preference behaviour is driven by colour and not intensity. (a**,**b)** Colour preference with reduced intensity through blue filter. **a**, Average daily preference between green, red (dim) and blue colour light with 100%, 50%, 25% or 10% of maximum (572 μW/cm^2^) intensity through blue filter. The fraction of flies in green, blue and red zones shown in the corresponding colour. Standard deviation between multiple consecutive days is shown in a shaded band around each line. 18 or 24 flies were used for each experiment. Black and white bars indicate dark and light part of LD cycle. **b**, Average avoidance of blue light at different intensities. The avoidance index was calculated (see Methods) for each hour from ZT 2-10, then averaged. **(c-e)** Preference in a two-choice assay with green light of different intensities. **c**, Average daily preference between two intensities of green colour light with the second option being 1, 0.5, 0.25, 0.1, 0.04 of the intensity of the first. **d**, Average preference for green light of the higher intensity in the first and last 3 hours of the day. Red line with shaded zone shows preference and standard deviation for green in green vs. red assay. The abscissa shows the ratio of the two intensities in logarithmical scale. **e**, Change in preference for green of higher intensity during first and last 3 hours of the day (M and E) and at ZT6 (Midday).

**Extended Data Figure 6. Flies tend to stay near food in the middle of the day. a-f**, Examples of fly positions during ZT3.5 to ZT8.5. Flies tend to stay near food when a green or red filter is nearest (a, c, e, f). When a blue filter is nearest to the food (b, d), flies enter the blue zone only briefly to eat. Ordinates represent the coordinate along the tube length. Horizontal orange lines represent edges of the colour zones.

**Extended Data Figure 7. Locomotor activity of wild-type flies and flies with abolished vision, MD neurons and both. a**, Average activity of flies in tubes with colour filters (black line) and without filters (orange line) under 12/12 LD. Light source was set at 2000 lux for tubes without filters. This was comparable to the irradiance of light transmitted through the filters in the colour preference experiments (see Methods). The morning (M) and evening (E) bursts of activity are similar in the two experiments. However, the level of activity during the middle of the day is significantly lower in tubes with filters. **b**, Abolishing vision and disabling activity of md neurons drastically reduce response to external light but still allow flies to move within the tube. **c**, *TrpA1*^*1*^ flies exhibit well-defined M and E bursts. The activity peaks seen in *per*^*0*^ data are not anticipatory M and E peaks, but instead are startle responses to lights turning on/off. **a-c**, Average activity was calculated as the distance moved by individual flies between consecutive video frames (captured at 1 frame/sec) and binned over 10-minute intervals. Black and white bars indicate dark and light portions of LD cycle.

**Extended Data Figure 8. Colour preference in additional genotypes. a-f**, Average daily preference between green, dim (red) and blue light for tested genotypes. The fraction of flies in green, blue and red zones are shown in the corresponding colour. Standard deviation between multiple consecutive days is shown in a shaded band around each line. 18 or 24 flies were used for each experiment. Black and white bars indicate dark and light part of LD cycle. **g**,**h**, Table of all genotypes, measured in three choice assays (**g**) and in blue/red assays (**h**), with an average fraction of flies spending time in blue zones during ZT 2-10 and p-values from Fisher’s exact test.

**Extended Data Figure 9. Correlation between colour preference and activity for wild-type and *period* mutants. a**, Colour preferences and activities of CantonS, *per*^*S*^ and *per*^*L*^ flies under 9.5/9.5 LD (19h), 12/12 LD (24h) and 14.75/14.75 LD (29.5h) cycles. The fraction of flies in green, blue and red zones is shown in the corresponding colour. Two experiments were carried out, each with 18 or 24 flies of each genotype. Average activity of flies is shown with a black line. Activity was calculated from video recordings as the distance moved by fly between two consecutive frames, binned over 3 minutes intervals, then normalized. M and E represent morning and evening bursts in locomotor activity. **b**, Timing of the M (squares) and E (circles) activity bursts in relation to peaks in green colour preference (green diamonds), under three separate LD cycles. The green peaks always appear near the end of the M activity and the start of the E activity when M/E occur during the lights-on portion of LD. **a**,**b**, Grey shaded regions represent lights-off portions of LD cycles.

**Extended Data Figure 10. The order of filters does not affect colour preference. a**, Sketch of a typical order of filters around tubes, with a total of 6 possible combinations. **b**, Example of 18 actual tubes in an experiment. Each tube is about 6 cm long and 5 mm wide. **c**,**d**, An example of the randomized order of filters (**c**) and the resulting colour preference data (**d**). Patterns in colour preference with randomized filter order were identical (within error bars) to that for a typical arrangement of filters as sketched in **a**. This indicates that the position of colour filters on neighbouring tubes did not affect fly behaviour. **d**, Fraction of flies in green, blue and red zones is shown in the corresponding colour, and standard deviation between multiple consecutive days is shown in shaded bands. Black and white horizontal bars indicate dark and light parts of LD cycle.

