## Supplementary Materials for "Daytime Colour Preference in Drosophila Depends on the Circadian Clock and TRP Channels"

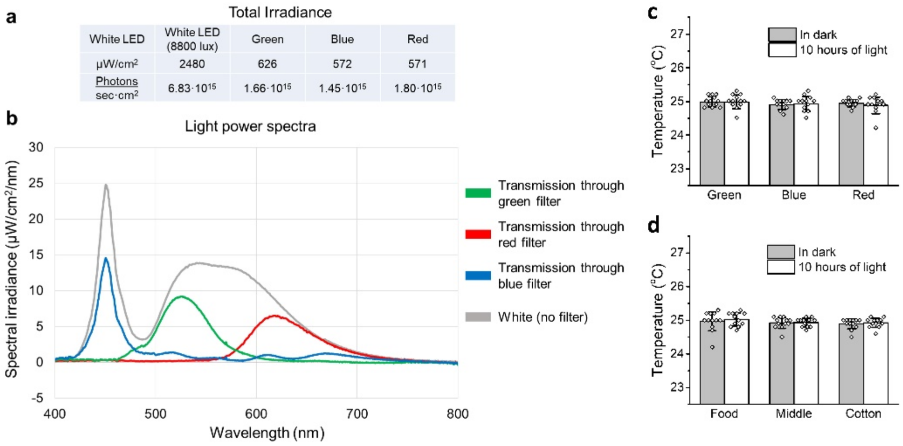


**Extended Data Figure 1. Properties of unfiltered and filtered light, and temperature inside the tubes.** **a**, Measured values for the broadband ‘white’ LED light of ~8800 lux used in the experiments and the corresponding total irradiance of the light before and after passing through the filters. **b**, Spectral irradiance of the LED light and light transmitted through the green, blue and red filters. Peaks for the filtered light are at 450 nm (blue), 528 nm (green), and 620 nm (red).**c,** Average temperature in the green, blue, and red zones, regardless of proximity to food or cotton plug. **d**, Average temperature near food, near cotton and in the middle, regardless of filter positions. **c,d,** Measurements were taken with lights turned off (black bars) and after lights have been on for 10 hours (white bars). Error bars are standard deviations from 12 independent measurements.

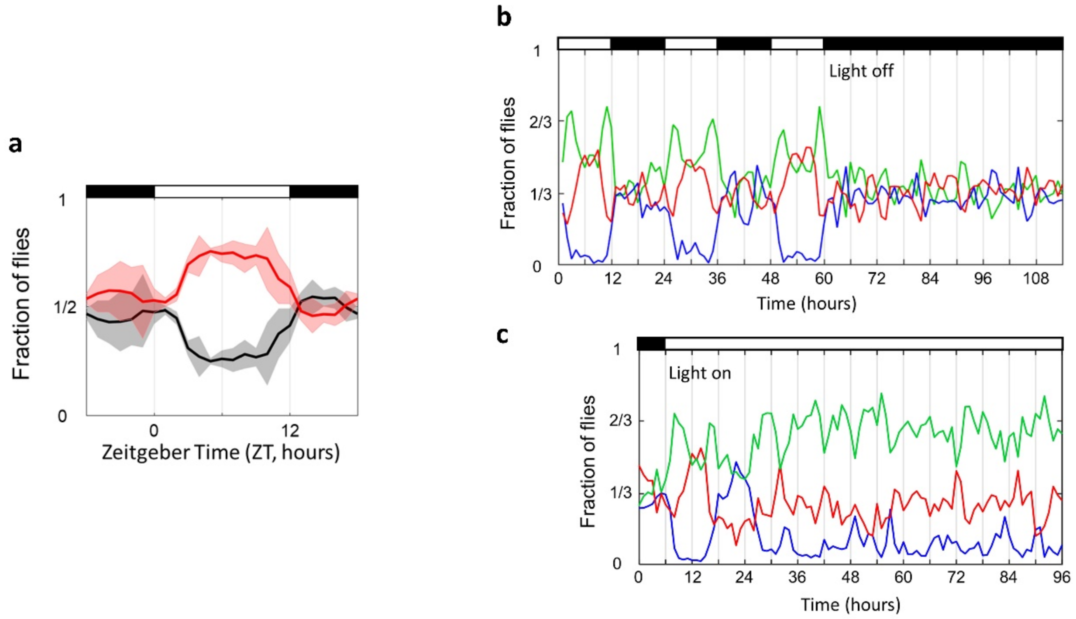


**Extended Data Figure 2. Flies can differentiate between dim light and dark, and lose preference pattern in constant conditions. a**, Average daily preference between red and dark zones for wild-type flies. The dark zone was obtained by overlaying three colour filters (Rosco #27, #382, #389). The fraction of flies in red and dark zones shown in red and black. Standard deviation between 4 consecutive days is shown in shaded bands around each line. 20 flies were used for the experiment. **b,c,** Distribution of flies across colour zones in 12:12 LD conditions, followed by constant darkness (**b**) or constant light (**c**) (N=24, each experiment). The fraction of flies in green, blue and red zones shown in the corresponding colour. **a,b**, Black and white horizontal bars indicate dark and light conditions.

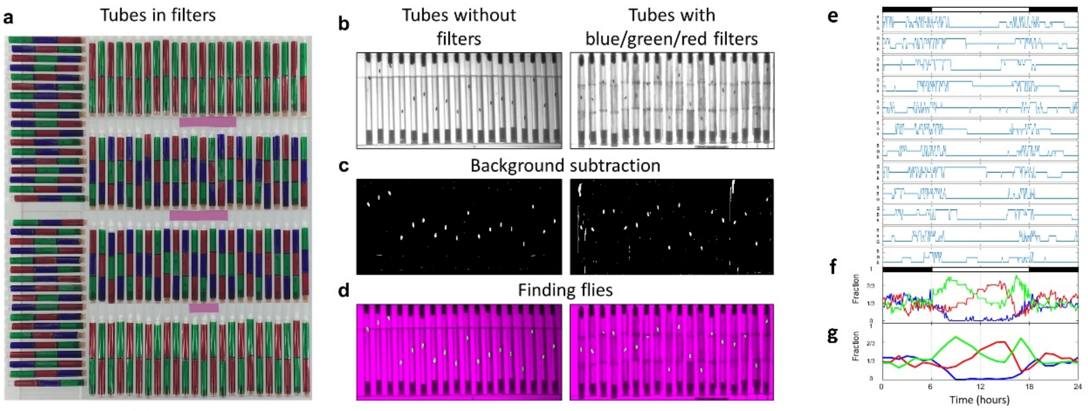


**Extended Data Figure 3. Experimental setup with tubes in filters. a,** Typical spatial arrangement of tubes (each approximately 6 cm long and 5 mm wide) in an experiment with three colour filters (green, blue, red) and two colour filters (green, red); N=132. **b**, Example frames of automatic tracking of flies in tubes with filters (right) and without filters (left). **c,** Background subtraction was applied after stabilizing video. **d,** Flies (green objects) were detected following digital filtering and noise elimination. **e**, Example of 12 randomly picked single fly trajectories, moving between green (G), blue (B) and red (R) zones over 24 hours. Fly positions were determined every minute but data shown are 5 minutes apart for clarity. **f**, Population-averaged colour preference of the 12 flies. As with the individual data, the time interval here is 5 minutes. **g**, Same colour preference but averaged over 1-hour intervals.

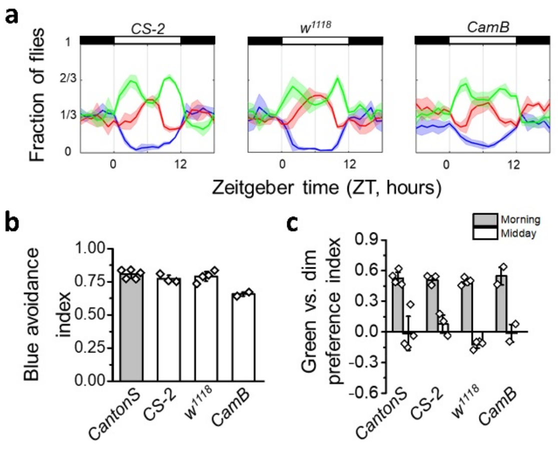


**Extended Data Figure 4. Preferences for colour in additional wild-type genotypes. a**, Preference among three zones for each genotype during one day. The fraction of flies in green, blue and red (dim) zones shown in the corresponding colour. An average 24 hour data (solid lines) and associated standard deviation (s.d., shaded bands) were generated from 3 experiments, each conducted for 4-6 days, with 18 or 24 flies of each genotype in each experiment. Black and white bars indicate dark and light part of LD cycle. **b**, Comparison of blue avoidance. The avoidance index was calculated (see Methods) for each hour from ZT 2-10, then averaged. **c**, Preference between green and dim light near timing of the first green peak ZT 2 and midday ZT 6. Preference for green is shown as positive and preference for dim light as negative.

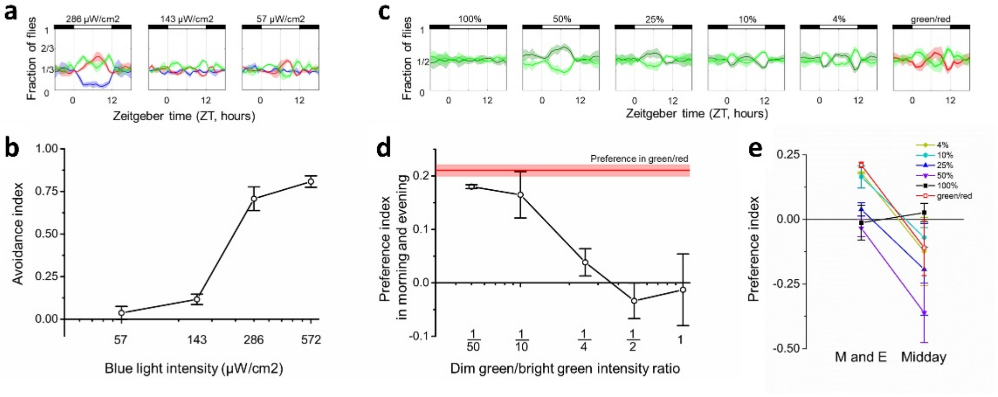


**Extended Data Figure 5. Colour preference behaviour is driven by colour and not intensity. (a,b)** Colour preference with reduced intensity through blue filter. **a**, Average daily preference between green, red (dim) and blue colour light with 100%, 50%, 25% or 10% of maximum (572 µW/cm$^2$) intensity through blue filter. The fraction of flies in green, blue and red zones shown in the corresponding colour. Standard deviation between multiple consecutive days is shown in a shaded band around each line. 18 or 24 flies were used for each experiment. Black and white bars indicate dark and light part of LD cycle. **b**, Average avoidance of blue light at different intensities. The avoidance index was calculated (see Methods) for each hour from ZT 2-10, then averaged. **(c-e)** Preference in a two-choice assay with green light of different intensities. **c**, Average daily preference between two intensities of green colour light with the second option being 1, 0.5, 0.25, 0.1, 0.04 of the intensity of the first. **d**, Average preference for green light of the higher intensity in the first and last 3 hours of the day. Red line with shaded zone shows preference and standard deviation for green in green vs. red assay. The abscissa shows the ratio of the two intensities in logarithmical scale. **e**, Change in preference for green of higher intensity during first and last 3 hours of the day (M and E) and at ZT6 (Midday).

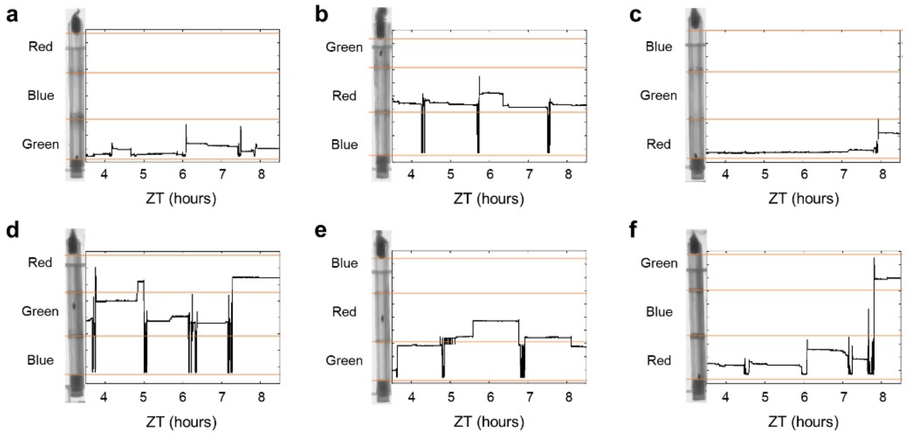


**Extended Data Figure 6. Flies tend to stay near food in the middle of the day. a-f**, Examples of fly positions during ZT3.5 to ZT8.5. Flies tend to stay near food when a green or red filter is nearest (a, c, e, f). When a blue filter is nearest to the food (b, d), flies enter the blue zone only briefly to eat. Ordinates represent the coordinate along the tube length. Horizontal orange lines represent edges of the colour zones.

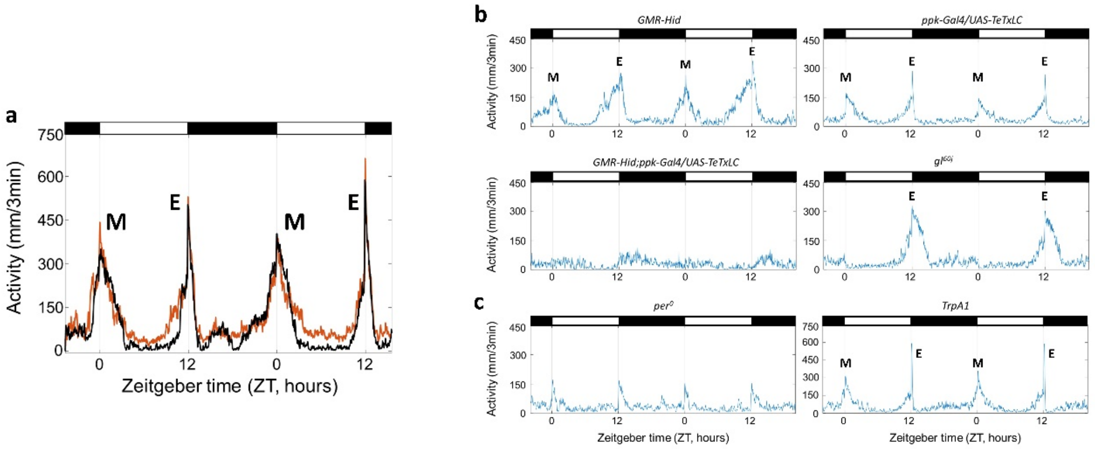


**Extended Data Figure 7. Locomotor activity of wild-type flies and flies with abolished vision, MD neurons and both. a**, Average activity of flies in tubes with colour filters (black line) and without filters (orange line) under 12/12 LD. Light source was set at 2000 lux for tubes without filters. This was comparable to the irradiance of light transmitted through the filters in the colour preference experiments (see Methods). The morning (M) and evening (E) bursts of activity are similar in the two experiments. However, the level of activity during the middle of the day is significantly lower in tubes with filters. **b**, Abolishing vision and disabling activity of md neurons drastically reduce response to external light but still allow flies to move within the tube. **c**, *TrpA1*$^{1}$ flies show well-defined M and E bursts. The activity peaks seen in *per*$^{0}$ data are not anticipatory M and E peaks, but instead are startle responses to lights turning on/off. **a-c,** Average activity was calculated as the distance moved by individual flies between consecutive video frames (captured at 1 frame/sec) and binned over 10-minute intervals. Black and white bars indicate dark and light portions of LD cycle.

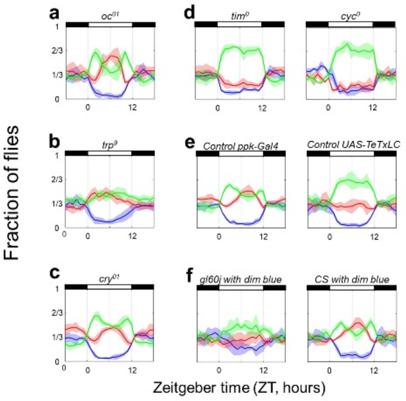


**g**

| Genotype | Total N | Avg % in blue | p-value |
|---|---|---|---|
| *CantonS* | 150 | 7% | |
| $gl^{60j}$ | 54 | 9% | 0.5486 |
| $norpA^{P24}$ | 54 | 9% | 0.5486 |
| *GMR-hid* | 48 | 8% | 0.7475 |
| $ninaE^{17}$ | 54 | 13% | 0.1595 |
| $Rh5^{2}$ | 48 | 17% | 0.0456 |
| $Rh6^{1}$ | 60 | 13% | 0.1695 |
| $oc^{01}$ | 42 | 9% | 0.5110 |
| $TrpA1^{1}$ | 66 | 8% | 0.7781 |
| $pain^{pc}$ | 54 | 26% | 0.0004 |
| *ppk > TeTxLC* | 84 | 19% | 0.0080 |
| *GMR-hid,ppk > TeTxLC* | 30 | 21% | 0.0308 |

| Genotype | Total N | Avg % in blue | p-value |
|---|---|---|---|
| $cry^{01}$ | 60 | 5% | 0.7618 |
| $trp^{9}$ | 54 | 11% | 0.3747 |
| *trpl* | 60 | 6% | 0.7618 |
| *gr28b* | 36 | 3% | 0.6941 |
| *ppk-Gal4* | 60 | 8% | 0.7675 |
| *UAS-TeTxLC* | 60 | 7% | 1.0000 |
| *CS-2* | 48 | 7% | 0.7476 |
| *CamB* | 36 | 14% | 0.1734 |
| *Gl60j with dim blue* | 24 | 23% | 0.0377 |
| *CS with dim blue* | 24 | 10% | 0.6725 |
| $pyx^{3}$ | 42 | 11% | 0.2427 |

**h**

| Genotype | Total N | Avg % in blue | p-value |
|---|---|---|---|
| *CantonS* | 76 | 14% | |
| $gl^{60j}$ | 46 | 13% | 1.0000 |
| $norpA^{P24}$ | 44 | 9% | 0.5683 |
| $oc^{01}$ | 34 | 18% | 0.7764 |
| $ninaE^{17}$ | 44 | 14% | 1.0000 |
| $Rh5^{2}$ | 38 | 32% | 0.0231 |

| Genotype | Total N | Avg % in blue | p-value |
|---|---|---|---|
| $Rh6^{1}$ | 52 | 25% | 0.1679 |
| $TrpA1^{1}$ | 58 | 16% | 1.0000 |
| $pyx^{3}$ | 36 | 15% | 1.0000 |
| $pain^{pc}$ | 38 | 45% | 0.0009 |
| *ppk > TeTxLC* | 42 | 31% | 0.0369 |

**Extended Data Figure 8. Colour preference in additional genotypes. a-f**, Average daily preference between green, dim (red) and blue light for tested genotypes. The fraction of flies in green, blue and red zones are shown in the corresponding colour. Standard deviation between multiple consecutive days is shown in a shaded band around each line. 18 or 24 flies were used for each experiment. Black and white bars indicate dark and light part of LD cycle. **g,h**, Table of all genotypes, measured in three choice assays (**g**) and in blue/red assays (**h**), with an average fraction of flies spending time in blue zones during ZT 2-10 and p-values from Fisher's exact test.

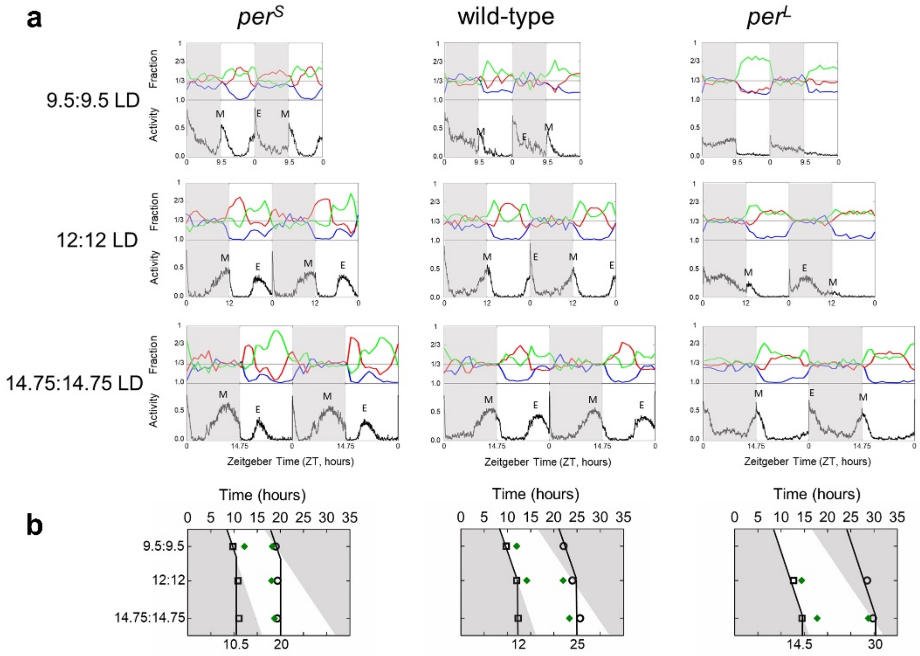


**Extended Data Figure 9. Correlation between colour preference and activity for wild-type and *period* mutants. a**, Colour preferences and activities of CantonS, *per*$^S$ and *per*$^L$ flies under 9.5/9.5 LD (19h), 12/12 LD (24h) and 14.75/14.75 LD (29.5h) cycles. The fraction of flies in green, blue and red zones is shown in the corresponding colour. Two experiments were carried out, each with 18 or 24 flies of each genotype. Average activity of flies is shown with a black line. Activity was calculated from video recordings as the distance moved by fly between two consecutive frames, binned over 3 minutes intervals, then normalized. M and E represent morning and evening bursts in locomotor activity. **b**, Timing of the M (squares) and E (circles) activity bursts in relation to peaks in green colour preference (green diamonds), under three separate LD cycles. The green peaks always appear near the end of the M activity and the start of the E activity when M/E occur during the lights-on portion of LD. **a,b**, Grey shaded regions represent lights-off portions of LD cycles.

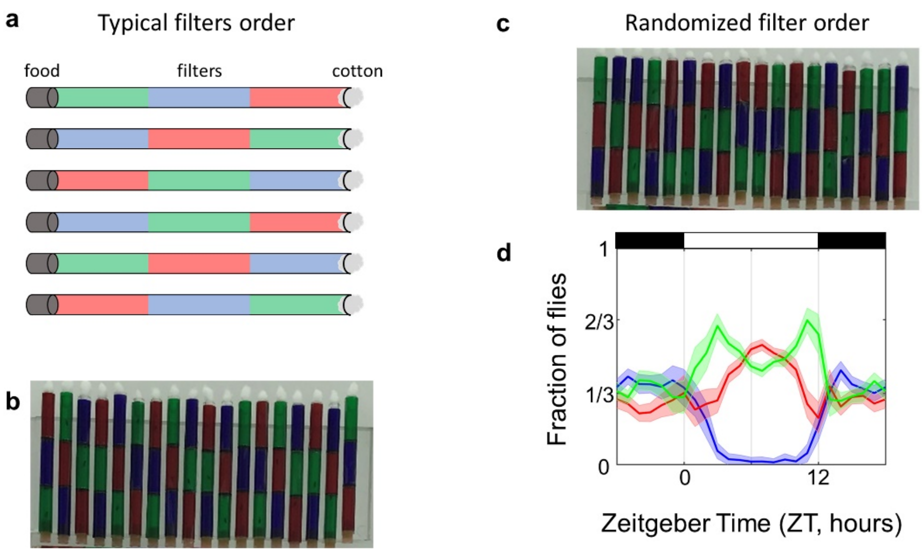


**Extended Data Figure 10. The order of filters does not affect colour preference. a**, Sketch of a typical order of filters around tubes, with a total of 6 possible combinations. **b**, Example of 18 actual tubes in an experiment. Each tube is about 6 cm long and 5 mm wide. **c**,**d,** An example of the randomized order of filters (**c**) and the resulting colour preference data (**d**). Patterns in colour preference with randomized filter order were identical (within error bars) to that for a typical arrangement of filters as sketched in **a**. This indicates that the position of colour filters on neighbouring tubes did not affect fly behaviour. **d**, Fraction of flies in green, blue and red zones is shown in the corresponding colour, and standard deviation between multiple consecutive days is shown in shaded bands. Black and white horizontal bars indicate dark and light parts of LD cycle.
